# MLL1 is required for maintenance of intestinal stem cells and the expression of the cell adhesion molecule JAML

**DOI:** 10.1101/2020.11.19.389528

**Authors:** Neha Goveas, Claudia Waskow, Kathrin Arndt, Julian Heuberger, Qinyu Zhang, Dimitra Alexopoulou, Andreas Dahl, Walter Birchmeier, Konstantinos Anastassiadis, A. Francis Stewart, Andrea Kranz

## Abstract

Epigenetic control is crucial for lineage-specific gene expression that creates cellular identity during mammalian development and in adult organism. Histone 3 lysine 4 methylation (H3K4) is a universal epigenetic mark. Mixed lineage leukemia (MLL1) is the founding member of the mammalian family of H3K4 methyltransferases. It was originally discovered as the main gene mutated in early onset leukemias and then found to be required for hematopoietic stem cell development and maintenance. However, the roles of MLL1 in non-hematopoietic tissues remain largely unexplored. To bypass hematopoietic lethality, we used bone marrow transplantation and conditional mutagenesis to discover that the most overt phenotype in *Mll1*-mutant mice is intestinal failure. Loss of MLL1 is accompanied by a differentiation bias towards the secretory lineage with increased numbers of goblet cells. MLL1 is expressed in intestinal stem cells (ISCs) and transit amplifying (TA) cells but at reduced levels in Paneth cells and not in the villus. MLL1 is required for the maintenance of intestinal stem cells (ISCs) and proliferation in the crypt. Transcriptome analysis implicate MLL1-dependent expression in ISCs of several transcription factors including *Pitx2, Gata4, Foxa1* and *Onecut2*, and also a cell adhesion molecule, *Jaml*. Reactive transcriptome changes in Paneth cells and organoids imply that JAML plays a key role in the crypt stem cell niche. All known postnatal functions of MLL1 relate to stem cell maintenance and lineage decisions thereby highlighting the suggestion that MLL1 is a master stem cell regulator.

**Author Summary:** The ability of adult stem cells to produce functional progenies through differentiation is critical to maintain function and integrity of organs. A fundamental challenge is to identify factors that control the transition from self-renewal to the differentiated state. Epigenetic factors amongst others can fullfill such a role. Methylation of histone 3 on lysine 4 (H3K4) is a posttranslational epigenetic modification that is associated with actively transcribed genes. In mammals, this epigenetic mark is catalyzed by one of six H3K4 methyltransferases, including the founding member of the family, MLL1. MLL1 is important for the precise functioning of the hematopoietic stem cell compartment. This raises the possibility of similar functions in other adult stem cell compartments. Due to its intense self-renewal kinetics and its simple repetitive architecture, the intestinal epithelium serves as a prime model for studying adult stem cells. We demonstrate that MLL1 controls intestinal stem cell proliferation and differentiation. Additionally, transcriptome analysis suggests a pertubation in the close interaction between intestinal stem cells and neighbouring Paneth cells through loss of junction adhesion molecule like (JAML). Our work sheds new light on the function of MLL1 for the control of intestinal stem cell identity.

## Introduction

Stem cells are cornerstones of tissue biology, ensuring homeostasis and regeneration in many organs, including epithelial tissues such as skin, intestine and mammary gland [1]. Stem cells are characterized by multipotency, which is the ability to differentiate into a restricted number of defined cell types, and self-renewal, which is the capacity to undergo infinite replicative cycles without losing stem cell identity [2]. The remarkable capacities of stem cells, particularly the restricted specificities of multipotency, rely on interplays between specific transcription factors and distinct epigenetic landscapes. Whereas the transcription factors involved in stem cell maintenance and differentiation have been clearly defined, epigenetic contributions are proving more elusive. For example, the transcription factor hierarchies in the stem cell paradigm, hematopoiesis, have been elegantly dissected [3]. However the contributions of DNA and histone methyltransferases to hematopoiesis are still emerging and indicate both specificities and the deeper complexities of epigenetic regulation[4–8].

Methylation of histone 3 on lysine 4 (H3K4) is one of the most conserved and widespread epigenetic systems [9]. H3K4 is methylated in euchromatic regions, with trimethylated H3K4 (H3K4me3) on nucleosomes surrounding active promoters, H3K4me2 marking transcribed regions and H3K4me1 relating to enhancers and active chromatin in general [10–14]. Mammals have six Set1/Trithorax-related methyltransferases that are encoded by three pairs of paralogous sister genes namely, *Mll1* (*Kmt2a*) and *Mll2* (*Kmt2b*), *Mll3* (*Kmt2c*) and *Mll4* (*Kmt2d*), *Setd1a* (*Kmt2f*) and *Setd1b* (*Kmt2g*). Each of the six methyltransferases reside in their own, large, protein complex. However all six complexes are based on a four membered scaffold termed WRAD for the subunits WDR5, RBBP5, ASH2L and DPY30 [15]. Functional differences between the six complexes potentially arise from the presence of additional subunits, which are usually shared by paralogous pairs or sometimes uniquely found in one of the six complexes.

Mixed lineage leukemia (*MLL1*) was the first mammalian gene identified as a Trithorax homologue and subsequently found to encode a mammalian Set1/Trithorax-type H3K4 methyltransferase (HMT) [16, 17]. In mice, MLL1 is first required at embryonic day 12.5 (E12.5) for definitive hematopoiesis [18, 19] and also required for the maintenance of adult hematopoietic stem cells (HSCs) [20, 21]. *MLL1*, but not its paralogue, *MLL2*, is a proto-oncogene because it can be activated by chromosomal translocations to promote leukemias without additional mutagenesis [22, 23]. Over 80 translocation partners have been identified including AF6 and AF9 [24]. Notably, *MLL1-AF6* and *-AF9* leukemias rely on *Mll2* expression [6]. Mouse studies also indicated that *MLL1-AF9* leukemiogenesis is entirely conveyed by overexpression of *Hoxa9* [25]. Conditional mutagenesis has also revealed MLL1 functions in satellite cells [26] and postnatal neural stem cells (NSCs) [27]. These observations raise the possibility that MLL1 regulates specific functions in stem cell compartments.

Due to its high turnover and hierarchical architecture, intestinal stem cells (ISCs) in the intestinal epithelium have become an adult stem cell paradigm. ISCs have been identified as either actively cycling crypt base columnar cells (CBCs) or quiescent label-retaining cells (LRCs) located at the +4 position from the crypt base [28]. Leucine-rich repeat-containing G protein-coupled receptor 5 (LGR5) is one of the best-characterized markers for the CBC class ISCs [29]. They generate transit amplifying (TA) daughter cells that give rise to the terminally differentiated progenies: absorptive enterocytes, secretory goblet, enteroendocrine and Paneth cells [30]. Except for Paneth cells, these cell types take 3 to 5 days to migrate up the villi and are shed into the intestinal lumen. Paneth cells reside at the base of the crypts in close association with ISCs and turn over at a slower rate [30].

The quiescent LRCs are active only during stress or injury. They represent a stem cell reserve to replace damaged ISCs [31]. Enterocytes, preterminal enteroendocrine cells, goblet cell precursors and Dll1^+^ secretory progenitors are also notable for their plasticity and can serve as a reservoir for lost stem cells [32, 33] [34]. Mature Paneth cells also show an injury-activated conversion to a stem cell like state [35]. Together with new perceptions in hematopoiesis [4, 6, 36], the dynamic plasticity of the intestinal crypt has expanded the stem cell paradigm [37], especially during replenishment after damage or inflammation.

To examine whether MLL1 plays additional roles in the adult, we employed ligand-induced conditional mutagenesis using a tamoxifen-inducible *Rosa26-CreERT2* line (*RC*) for near-ubiquitous Cre recombination to discover that MLL1 is required for the maintenance of the ISC compartment and the balance between secretory and absorptive cell lineages in the adult intestine.

## Results

### Intestinal functions collapse after loss of MLL1 in adult mice

To explore MLL1 functions, we utilized a multipurpose allele that can be converted from one state to another using FLP and Cre recombination (S1A-S1C Figs) [38, 39]. Homozygous embryos carrying the targeted allele, *Mll1*^*A/A*^, developed normally until E12.5 when they displayed pallor of the liver and were smaller (S1D Fig). After E13 no live *Mll1*^*A/A*^ embryos were found (S1 Table). After FLP and Cre recombination, *Mll1*^*FC/FC*^ embryos displayed the same phenotype indicating that both A and FC are true null alleles in concordance with the loss of MLL1 protein [40]. Furthermore the *Mll1*^*A/A*^ and *Mll1*^*FC/FC*^ phenotype recapitulated another likely null allele [6] with all three homozygous mice presenting the same embryonic lethality due to the failure to engage definitive hematopoiesis [20].

To identify postnatal roles of MLL1, conditional mutagenesis using *Rosa26-CreERT2* was applied to 2 month old adults. As expected, mice lacking MLL1 developed severe bone marrow cytopenia and died or had to be sacrificed on average within two weeks (Figs 1A and 1B). As previously reported using the same conditional allele, *in vitro* deletion of *Mll1* in KSL-enriched HSCs from *Mll1*^*F/F; RC*/+^ mice resulted in significant downregulation of *Hoxa9*, *Meis1*, *Mecom/Evi1* and *Prdm16* [6].

**Fig 1.**
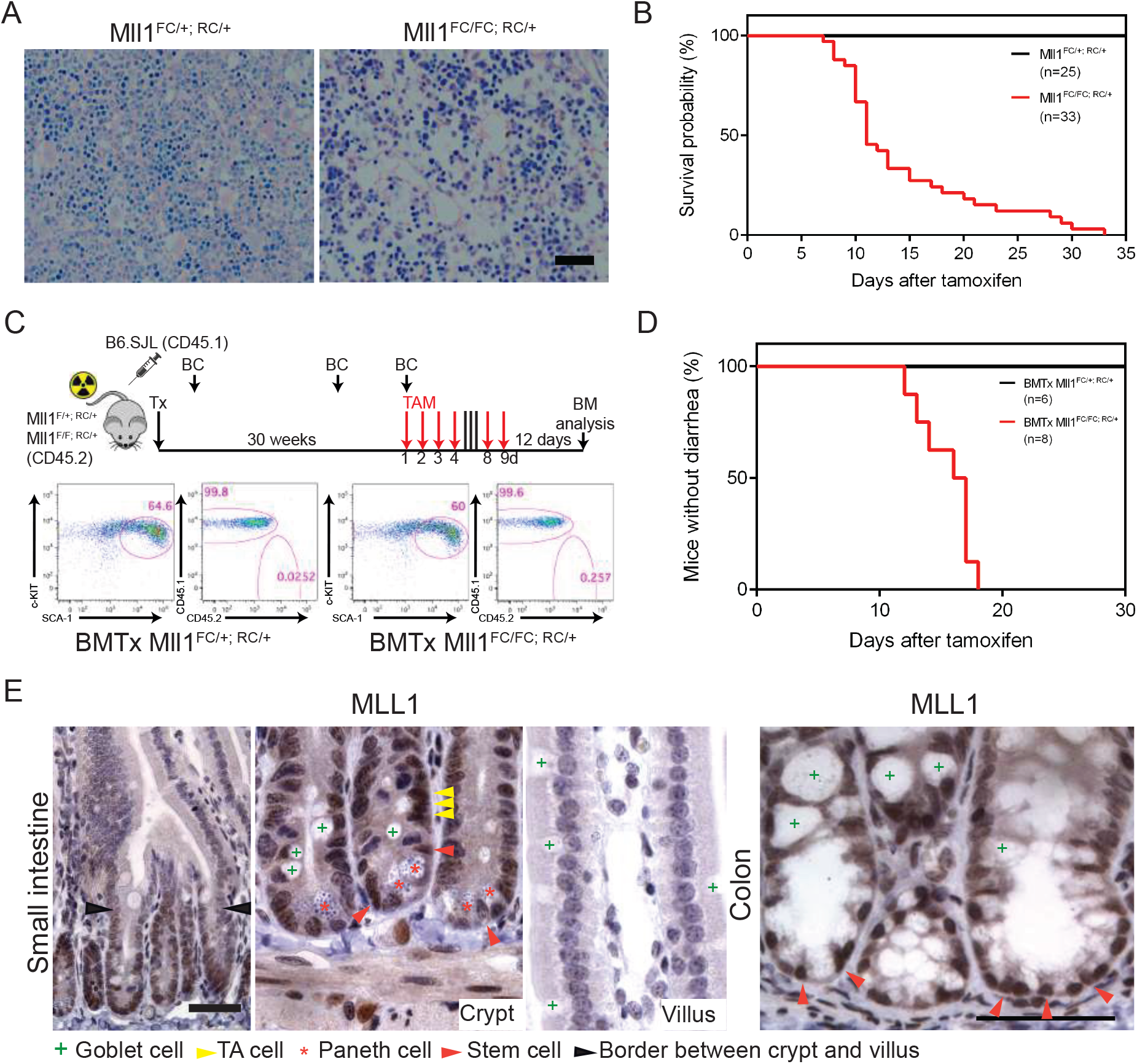
Loss of MLL1 in adult mice. (A) Giemsa-stained sections from the femur of *Mll1*^*FC*/+; *RC*/+^ and *Mll1*^*FC/FC; RC*/+^ mice. Two weeks after the last tamoxifen gavage mice were sacrificed and the femurs were dissected, decalcified, sectioned and stained. Bone marrow cellularity was severely decreased in *Mll1*^*FC/FC; RC*/+^ mice. Scale bar 100 μm. (B) Kaplan-Meier survival curve. The first day of tamoxifen gavage was day zero. All *Mll1*^*FC*/+; *RC*/+^ mice (n=25) survived whereas all *Mll1*^*FC/FC; RC*/+^ mice (n=33) died within 33 days after tamoxifen induction with a median survival of 11 days. (C) Scheme of the experimental setup for bone marrow transplantation. Donor bone marrow from B6.SJL (CD45.1^+^) mice was transplanted into lethally irradiated *Mll1*^*F*/+; *RC*/+^ and *Mll1*^*F/F; RC*/+^ recipients (CD45.2^+^). Blood chimerism (BC) was measured three times. After 30 weeks *Mll1* deletion was achieved by administrating tamoxifen (TAM). FACS analysis for KSL-Slam enriched HSCs (Kit^+^ Sca1^+^ Lin^−^ CD48^−^ CD150^+^ CD34^−^ CD135^−^) showed comparable numbers in BMTx *Mll1*^*FC*/+; *RC*/+^ and *Mll1*^*FC/FC; RC*/+^ mice. Dot plots show Lin^−^ CD48^−^ CD150^+^ CD34^−^ CD135^−^ gated bone marrow (BM) cells of indicated genotypes resolved for the expression of c-Kit and Sca-1. Donor and host cells are distinguished by surface markers CD45.1 and CD45.2. (D) Kaplan-Meier analysis for the onset of diarrhea. Tamoxifen was given by gavage for 6 days to *Mll1*^*F*/+; *RC*/+^ (n=6) and *Mll1*^*F/F; RC*/+^ (n=8). The first day of tamoxifen gavage was day zero. While all BMTx mice with the genotype *Mll1*^*FC*/+; *RC*/+^ remained healthy, all BMTx *Mll1*^*FC/FC; RC*/+^ mice developed diarrhea with a median of 16.5 days. (E) Antibody staining (brown) showed that MLL1 is expressed in crypts of the small and large intestine but absent in the villus (hematoxylin, purple). Scale bars are 50 μm.

To bypass the bone marrow related lethality and thereby uncover non-hematopoietic phenotypes, bone marrow from wild type (wt) B6.SJL mice was transplanted into lethally irradiated *Mll1*^*F*/+; *RC*/+^ or *Mll1*^*F/F; RC*/+^ mice. After stable engraftment tamoxifen gavage induced widespread Cre-mediated excision of *Mll1* (Fig 1C). Examination of the bone marrow confirmed the successful and near-complete reconstitution of the hematopoietic stem cell compartment by wt donor cells. FACS analysis for KSL-Slam enriched hematopoietic stem cells showed comparable frequencies in bone marrow transplanted (BMTx) *Mll1*^*FC*/+; *RC*/+^ and *Mll1*^*FC/FC; RC*/+^ mice with the hematopoietic compartment comprised only of wt cells of CD45.1 origin (Fig 1C). Notably, the BMTx *Mll1*^*FC/FC; RC*/+^ mice suffered from diarrhea and wasting (Fig 1D). These data indicate that MLL1 is not only required in the hematopoietic compartment but also elsewhere and this additional requirement is similarly critical for survival as the hematopoietic requirement.

The small and large intestine harbor their stem cell compartments at the base of the crypt. MLL1 is strongly expressed at the crypt bottom and in the TA compartment whereas it is absent in differentiated cells above the TA compartment (Figs 1E and S1E). RNA profiling of sorted ISCs and Paneth cells confirmed expression of *Mll1* and the other family members in both cell types however more strongly in ISCs (S1F Fig). In the mutant small intestine of BMTx *Mll1*^*FC/FC; RC*/+^ mice, expression of MLL1 was efficiently ablated (Fig 2A). Loss of stem cell markers olfactomedin (OLFM4) and SOX9 suggested depletion of ISCs in the small intestine of BMTx *Mll1*^*FC/FC; RC*/+^ mice (Fig 2A). Consistent with this, the mutant showed decreased proliferation in the crypt as demonstrated by strongly reduced expression of the mitotic marker Ki67 (Fig 2A). However, the intestinal epithelium of *Mll1*^*FC/FC; RC*/+^ mice revealed no apparent change in global H3K4 mono-, di- and trimethylation (S1G Fig).

**Fig 2.**
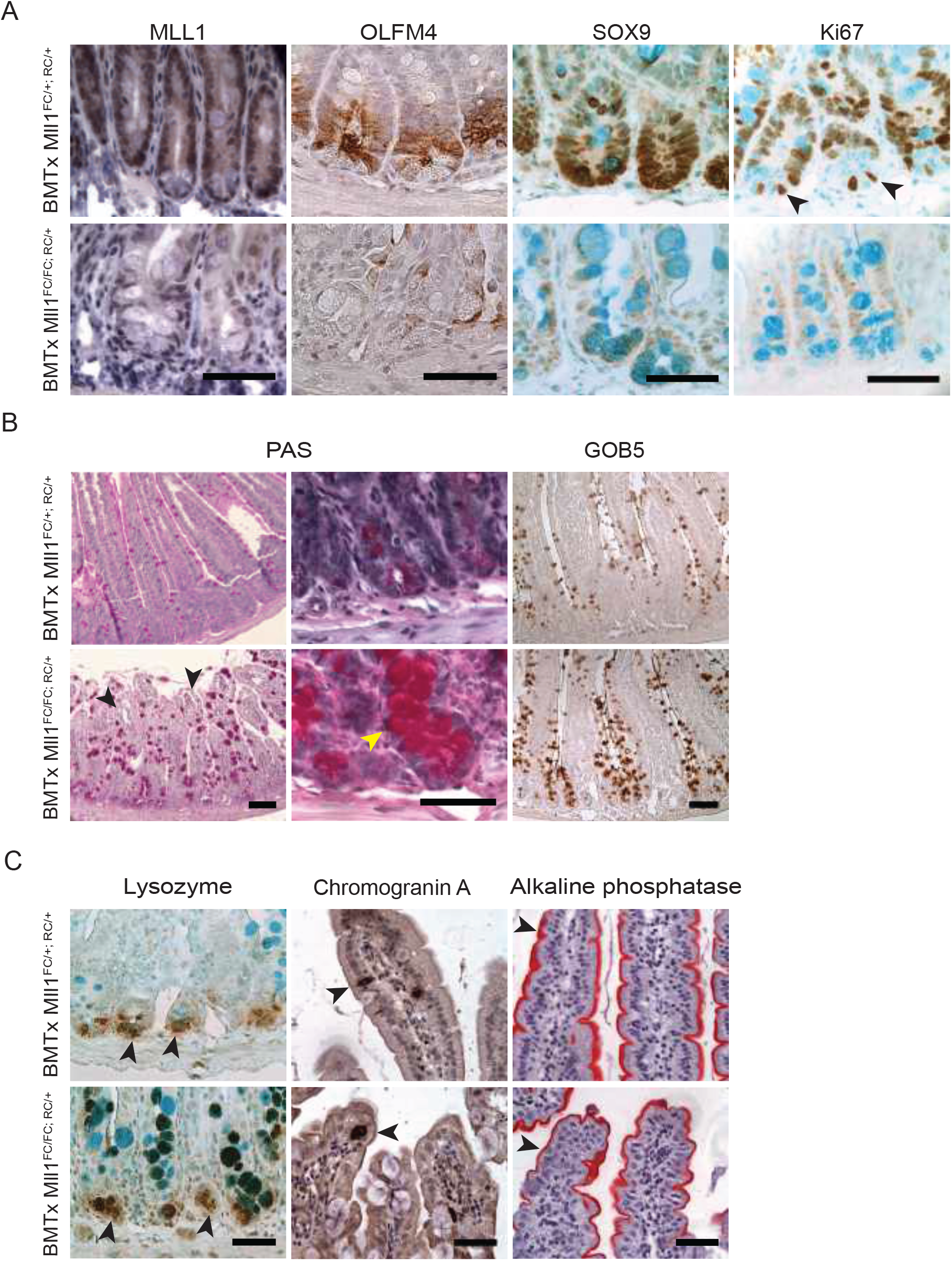
*Mll1* deletion in BMTx mice leads to loss of stem and proliferating cells and increased goblet cells. (A) Antibody stainings showed reduced MLL1, OLFM4 and SOX9 expression in BMTx *Mll1*^*FC/FC; RC*/+^ intestine compared to controls. Hematoxylin was used as a counterstain for MLL1 and OLFM4 immunohistochemistry (IHC). Expression of Ki67, a marker for proliferating cells was reduced in BMTx *Mll1*^*FC/FC; RC*/+^ mutant intestinal sections. Arrowheads point toward proliferating ISCs. Alcian blue was used as a counterstain for SOX9 and Ki67 IHC, which also revealed enlarged goblet cells (turquoise) in the crypts of BMTx *Mll1*^*FC/FC; RC*/+^ sections. Scale bars are 50 μm. (B) PAS stain to examine goblet cells in villi and crypts of BMTx intestine. Black arrowheads point towards vacuolar structures. Yellow arrowhead points to a mislocalized goblet cell in BMTx *Mll1*^*FC/FC; RC*/+^ crypt. Left panels scale bar 100 μm; middle panels scale bar 50 μm. GOB5 antibody staining of BMTx *Mll1*^*FC/FC; RC*/+^ intestinal sections. Right panels scale bar 100 μm. (C) Left panels; lysozyme antibody staining reveals that Paneth cell numbers remain unchanged. Arrowheads point at Paneth cells. Alcian blue was used as a counterstain and marks goblet cells (turquoise). Middle panels; chromogranin A antibody stain, arrowheads point to the sparse enteroendocrine cells (dark brown) in the villi. Right panels; red enterocytes covering the villi were visualized by alkaline phosphatase staining. Hematoxylin was used as a counterstain for chromogranin A IHC and alkaline phosphatase histochemical staining. Scale bars are 50 μm.

Shortened villi with distorted morphology including vacuolar structures at the tip indicated diminished replenishment of cells into the villus (Fig 2B). Furthermore we observed increased numbers of enlarged goblet cells distributed irregularly along the villus and also ectopically in the crypt (Fig 2B). However, Paneth cells appeared unchanged possibly due to their longer life span (Fig 2C). Similarly, as evaluated by chromogranin A and alkaline phosphatase, the enteroendocrine and absorptive lineages appeared to be unaffected (Fig 2C).

Without bone marrow rescue, *Mll1*^*FC/FC; RC*/+^ mice showed the same defects with depletion of ISCs, decreased proliferation and a distortion of the secretory lineage (S2A-S2C Figs). Differentiation into the enteroendocrine and absorptive lineage was also apparently unaffected (S2D Fig). Notably the hallmark of Wnt signalling, nuclear β-catenin, was also unaffected (S2E Fig).

### Intestinal epithelium-specific *Mll1* conditional mutagenesis recapitulates ubiquitous deletion

To delete *Mll1* exclusively in the adult intestine we employed the tamoxifen-inducible gut epithelium-specific *Villin-CreERT2* strain [41]. After tamoxifen administration *Mll1*^*FC/FC; Vil-CreERT2*/+^ mice lost weight compared to control mice (Fig 3A). In agreement with our observations after ubiquitous deletion of *Mll1*, OLFM4, SOX9 and proliferation were markedly decreased, goblet cells were increased whereas Paneth and enteroendocrine cell numbers were unchanged (Figs 3B and 3C).

**Fig 3.**
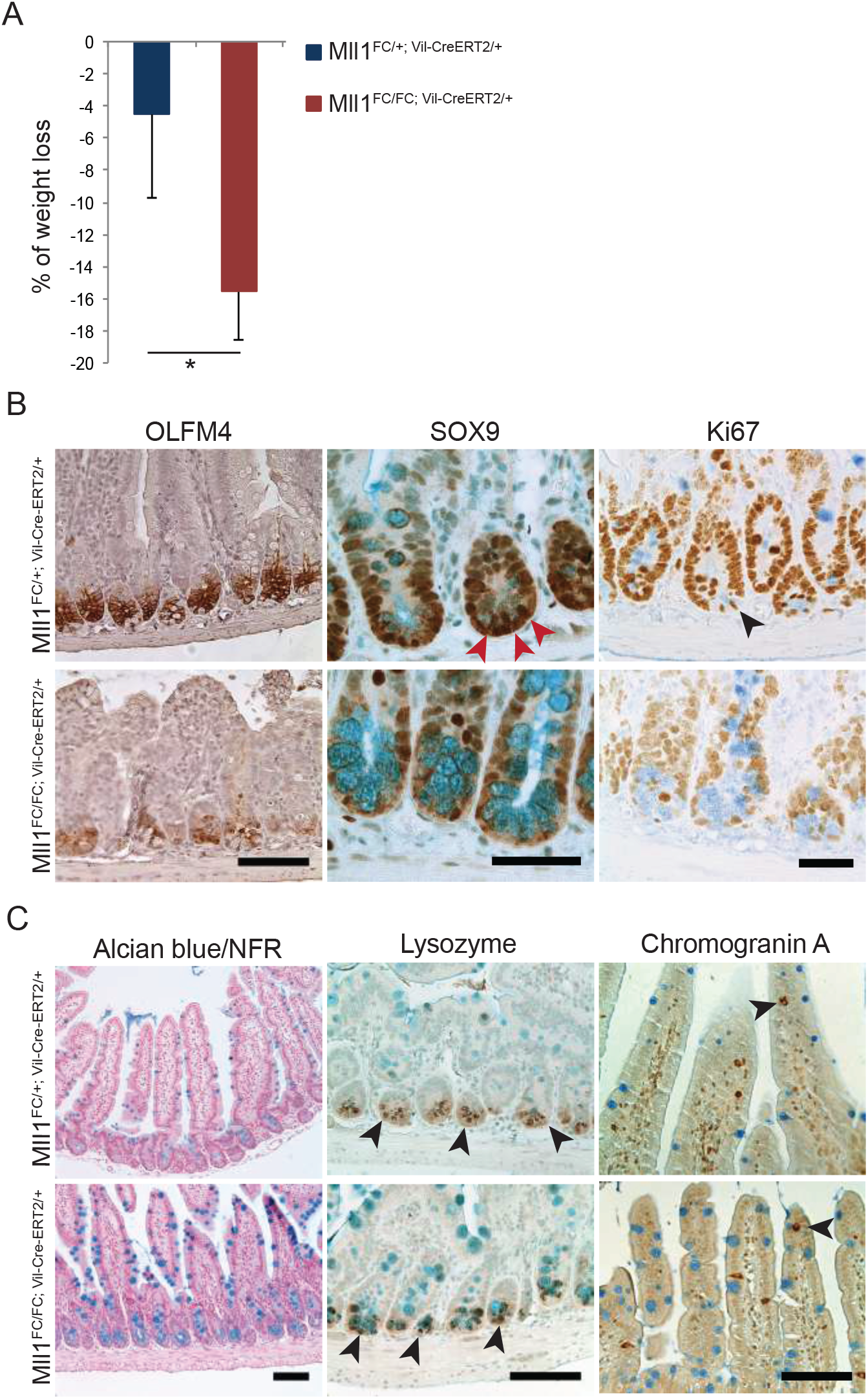
Decreased ISCs and increased goblet cells after intestinal specific loss of MLL1. (A) Percent of weight loss of *Mll1*^*FC*/+; *Vil-Cre-ERT2*/+^ (n=5) and of *Mll1*^*FC/FC; Vil-Cre-ERT2*/+^ (n=9) mice. Mean ± s.d. is shown (p=0.029, Student’s *t*-test). (B) Decrease in ISC markers, OLFM4 and SOX9 in *Mll1*^*FC/FC; Vil-Cre-ERT2*/+^ intestinal sections. SOX9^+^ ISCs in *Mll1*^*FC*/+; *Vil-Cre-ERT2*/+^ intestinal sections are marked with red arrowheads. Proliferating cells in the TA compartment as well as proliferative ISCs (arrowhead) are reduced in *Mll1*^*FC/FC; Vil-Cre-ERT2*/+^ sections. Alcian blue was used as a counterstain after staining for SOX9 and Ki67 and marks goblet cells (turquoise). Scale bars are 100 μm for OLFM4 and 50 μm for SOX9 and Ki67. (C) Left panels; alcian blue staining of goblet cells with nuclear fast red (NFR) to stain nuclei. Middle panels; Paneth cells visualized by staining of granules containing lysozyme (arrowheads). Right panels; enteroendocrine cells stained with chromogranin A antibody (brown; arrowheads) in the villi. Scale bars are 100 μm.

### Transcriptome analysis identifies key intestine specific transcription factors and *Jaml* as central to the crypt stem cell niche

For transcriptome analysis, ISCs and Paneth cells were isolated 4 and 10 days after tamoxifen administration from *Mll1*^*FC*/+; *Lgr5-eGFP-CreERT2*/+^ and *Mll1*^*FC/FC; Lgr5-eGFP-CreERT2*/+^ littermates (S3 Fig). Using 75 base-pair reads, 20-37 million reads per sample with high levels of uniqueness (70-77% in Lgr5^+^ ISCs and 60-76% in Paneth cells; S4A-S4C Figs) and comparable mappability (99%) were obtained. Principal component analysis (PCA) revealed that our datasets are in good agreement with published datasets obtained from sorted ISCs and Paneth cells [35, 42] (S4D Fig).

We applied DESeq2 to analyze differentially expressed genes (DEGs). For the 4 day Lgr5^+^ stem cell profile, only 87 and 49 genes were up- or downregulated at a 5% false discovery rate (FDR) after removal of MLL1 (Fig 4A, Supplementary excel file 1). However, none of the upregulated transcripts were increased by more than log2-fold and by DAVID analysis were mainly related to diverse terms such as ‘response to metal ion’, ‘organic acid metabolic process’ and ‘regulation of lipid metabolic process’ (Fig 4B). In contrast the most significant terms associated with the downregulated mRNAs were ‘regulation of gene expression’, ‘epithelial cell differentiation’ and ‘cell proliferation’ (Fig 4B). The 10 day profile revealed 179 DEGs, of which 105 were upregulated, 74 were downregulated with significant overlaps to the 4 day profile (Supplementary excel file 1, Table 1).

**Fig 4.**
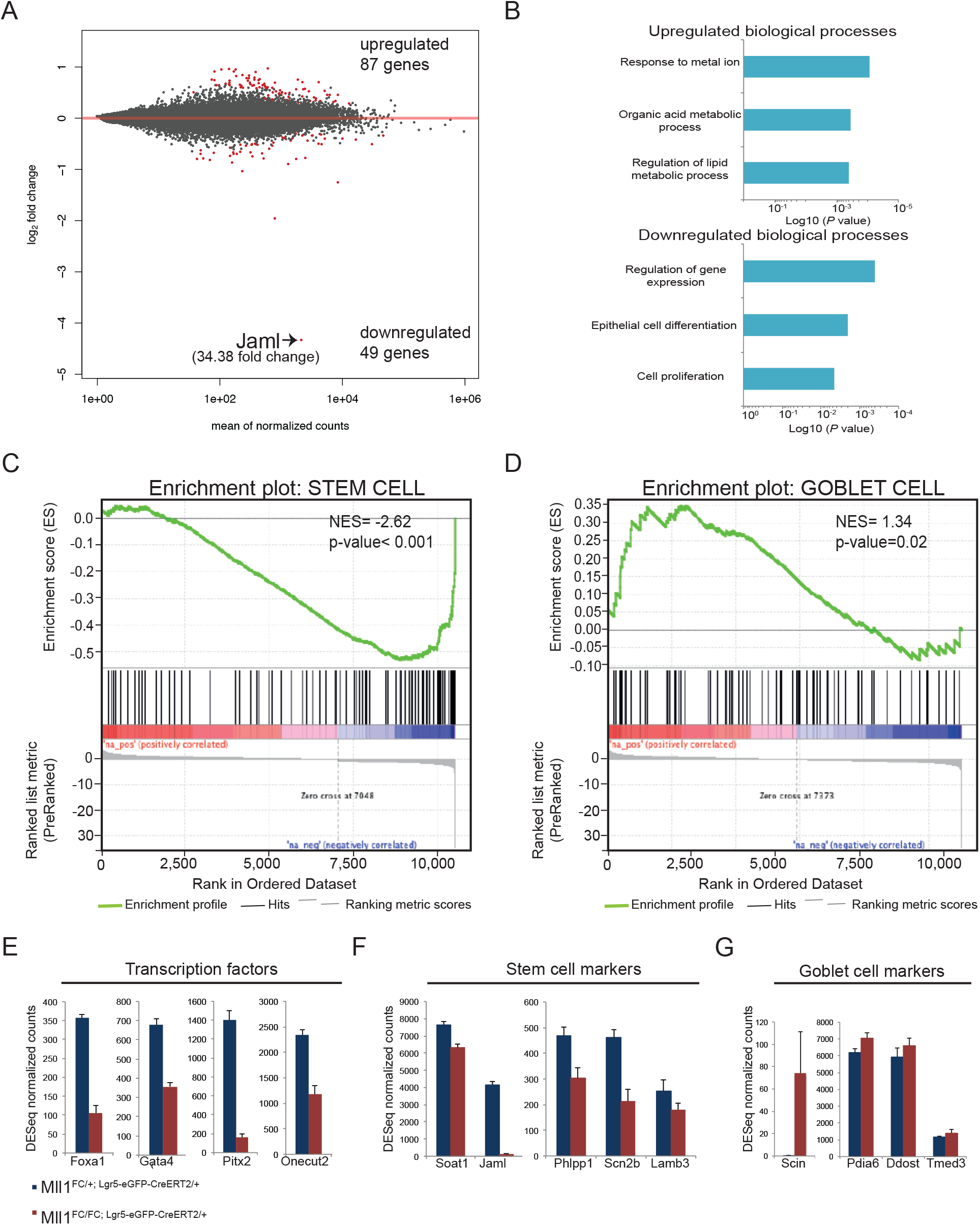
RNA profiling of *Mll1*^*FC/FC; Lgr5-eGFP-CreERT2*/+^ and control ISCs. (A) ISCs were sorted from control (*Mll1*^*FC*/+; *Lgr5-eGFP-CreERT2*/+^) (n=4) and *Mll1*^*FC/FC; Lgr5-eGFP-CreERT2*/+^ (n=4) mice 4 days after tamoxifen induction was completed and subjected to RNA profiling. MA plot visualizing the log2-fold change differences according to expression levels of ISCs. Red dots represent significant DEGs at a 5% FDR. *Jaml* is the top downregulated gene. (B) Plots show biological processes (BP) that are enriched in genes up- or downregulated in *Mll1*^*FC/FC; Lgr5-eGFP-CreERT2*/+^ compared to control ISCs. Analysis was performed using the gene ontology (GO)/BP/FAT database of DAVID 6.8. (C) (D) GSEA shows significant negative or positive correlation of genes from the stem (C) and goblet cell (D) signature gene set in *Mll1*^*FC/FC; Lgr5-eGFP-CreERT2*/+^ compared to control ISCs (4 days after tamoxifen). The signature gene sets originate from [44]. NES: normalized enrichment score. (E) DESeq normalized counts for genes coding for transcription factors downregulated in *Mll1*^*FC/FC; Lgr5-eGFP-CreERT2*/+^ compared to control ISCs (4 days after tamoxifen). Mean+s.d. is shown; n=4; p<0.05, Wald test. (F) DESeq normalized counts for genes coding for ISC markers downregulated in *Mll1*^*FC/FC; Lgr5-eGFP-CreERT2*/+^ compared to control ISCs (4 days after tamoxifen). Mean+s.d. is shown; n=4; p<0.05, Wald test (G) DESeq normalized counts for genes coding for goblet cell markers upregulated in *Mll1*^*FC/FC; Lgr5-eGFP-CreERT2*/+^ compared to control ISCs (4 days after tamoxifen). Mean+s.d. is shown; n=4; p<0.05, Wald test.

**Table 1.** Downregulated mRNAs common to both 4 and 10 day stem cell profiles.

| Gene |  | 4d C | KO | FC | 10d C | KO | FC |
| --- | --- | --- | --- | --- | --- | --- | --- |
| *Jaml/Amica1* | Junction Adhesion Molecule Like | 4124 | 119,9 | **34,4** | 3779 | 113,4 | **33,3** |
| *Pitx2* | Paired Like Homeodomain 2 | 1399,6 | 162,9 | **8,6** | 854,8 | 34,7 | **24,6** |
| *Far1* | FattyAcyl-CoA Reductase 1 | 402,3 | 229,1 | **1,8** | 231,8 | 47,4 | **4,9** |
| *Foxa1/Hnf3α* | Forkhead Box A1 | 356,8 | 105 | **3,4** | 428,4 | 93,9 | **4,6** |
| *Onecut2/Hnf6β* | Onecut Homeobox 2 | 2334,5 | 1177,1 | **2,0** | 2411,9 | 583 | **4,1** |
| *Gata4* | GATA Binding Protein 4 | 676,4 | 356,9 | **1,9** | 508,2 | 136,7 | **3,7** |
| *Ces2g* | Carboxyl esterase 2 | 275,2 | 177 | **1,6** | 209,9 | 58,6 | **3,6** |
| *Pla2g2a* | Phospholipase A2 group IIA | 291,3 | 119 | **2,5** | 215,7 | 68,2 | **3,2** |
| *Hspa8* | Hsp 70 family | 12104 | 4655,8 | **2,6** | 7654,5 | 25856 | **3,0** |
| *E230029C05* | ncRNA | 221,9 | 115,8 | **1,9** | 203,4 | 70,9 | **2,9** |
| *Casc4* | Cancer Susceptible 4 | 120,5 | 61,4 | **2,0** | 255,1 | 93,9 | **2,7** |
| *Anpep* | Alanyl amino peptidase N | 855,8 | 562,5 | **1,5** | 393,1 | 150,8 | **2,6** |
| *Defa29* | Defensinα 29 | 859,4 | 556,3 | **1,5** | 860,3 | 336,2 | **2,6** |
| *Lyz1* | Lysozyme | 10861 | 7181,7 | **1,5** | 8334,4 | 3429 | **2,4** |
| *Defa3* | Defensinα 3 | 342,6 | 239,5 | **1,4** | 521,2 | 214,7 | **2,4** |
| *Lcp1* | Lymphocyte Cytosolic P1 | 824,4 | 576 | **1,4** | 662 | 291 | **2,3** |
| *Defa35* | Defensinα 35 | 831,4 | 555,4 | **1,5** | 938,4 | 431,4 | **2,2** |
| *Gm10925* | pseudogene? | 18872 | 14115 | **1,3** | 30453 | 14252 | **2,1** |
| *Afap1l1* | Lnc-Afap1/1 | 485,5 | 354,5 | **1,4** | 420,6 | 197 | **2,1** |
| *Phlpp1* | PH Domain Leucine Rich Phosphatase | 468,7 | 302,8 | **1,6** | 480,5 | 227 | **2,1** |
| *Fap1* | Familial Adenomatous Polyposis 1 | 990,5 | 768,4 | **1,3** | 1459,8 | 690,9 | **2,1** |
| *Casp6* | Caspase 6 | 4727,6 | 3356,6 | **1,4** | 3861,6 | 1835 | **2,1** |
| *mt-Atp6* | Mitochondrial encoded ATP Synthase S6 | 49048 | 40226 | **1,2** | 61873 | 29894 | **2,1** |
| *Fam149a* | Toll-like receptor 3? | 170,7 | 97,6 | **1,8** | 193,9 | 94,3 | **2,1** |
| *Zfpm1* | FOG1 (Friend of Gata 1) | 198,5 | 146 | **1,4** | 216,6 | 106,2 | **2,0** |

mRNAs expressed more than 150 reads in wt and downregulated more than 2 fold in the 10 day profile are shown. FC, fold change.

Gene set enrichment analysis (GSEA) of both 4 and 10 day ISC profiles revealed that overall ISC signature genes were downregulated whereas goblet cell signature genes were upregulated (Figs 4C-4G; S5A and S5B Figs), which concords with our immunohistochemical analyses. We focused on the overlap between the 4 and 10 day downregulated mRNAs. The transcription factors *Pitx2, Foxa1*, *Onecut2* and *Gata4* are prominent (Table 1; Fig 4E). The expression of *Foxa1* and *Pitx2* and was evaluated by qRT-PCR and good agreement to the RNA-sequencing mRNA reads was found (S5C Fig). These data suggest a role for MLL1 in maintaining the transcriptional identity of ISCs.

However the top downregulated gene in both stem cell profiles (34.7 and 33.3 fold down) was junction adhesion molecule like (*Jaml* or *Amica1*; Figs 4A, 4F, S5C) previously identified as an ISC signature gene [43, 44]. It is highly expressed in Lgr5^+^ ISCs, where it appears to rely completely on MLL1, but not in Paneth cells (at least 55 fold lower expressed in Paneth cells; S5D Fig). JAML is a 65 kDa type I transmembrane glycoprotein in the JAM subset of the immunoglobulin superfamily. JAML mediates adhesion of monocytes to endothelial cells and neutrophil migration across epithelial cell monolayers through interaction with Coxsackie and adenovirus receptor (CXADR or CAR) in tight junctions [45]. However, the cognate receptor in the intestinal epithelium is unknown. Notably the transcript for CAR-like soluble protein (*Clsp*) (GM1123) is upregulated in both 4 and 10 day profiles (Supplementary excel file 1). CLSP is closely related to CXADR however it lacks a transmembrane domain [46].

### Loss of MLL1 in ISCs provokes transcriptional changes in Paneth cells

ISCs are anchored in the crypt in close association with Paneth cells. In order to elucidate whether the transcriptional changes in ISCs influenced the neighboring Paneth cells, we also analyzed Paneth cell transcriptional profiles 4 days after deletion of *Mll1* in Lgr5^+^ ISCs, with 198 and 72 transcripts up- and downregulated respectively (Fig 5A, Supplementary excel file 1). The most significant terms associated with downregulated mRNAs relate to perturbation of protein folding and homeostasis in the endoplasmic reticulum (Figs 5B-5D). In contrast upregulated mRNAs associate with metabolic changes. Strikingly, transcripts of genes belonging to all five of the respiratory chain complexes were upregulated (Figs 5B-5D). Paneth cells normally run on glycolysis with lactate as the end product whereas ISCs depend on mitochondrial oxidative phosphorylation [47]. These transcriptional changes suggest that loss of MLL1 in the stem cell compartment provokes changes in Paneth cells and indeed expression of Paneth cell marker genes was downregulated (Figs 5C and 5D).

**Fig 5.**
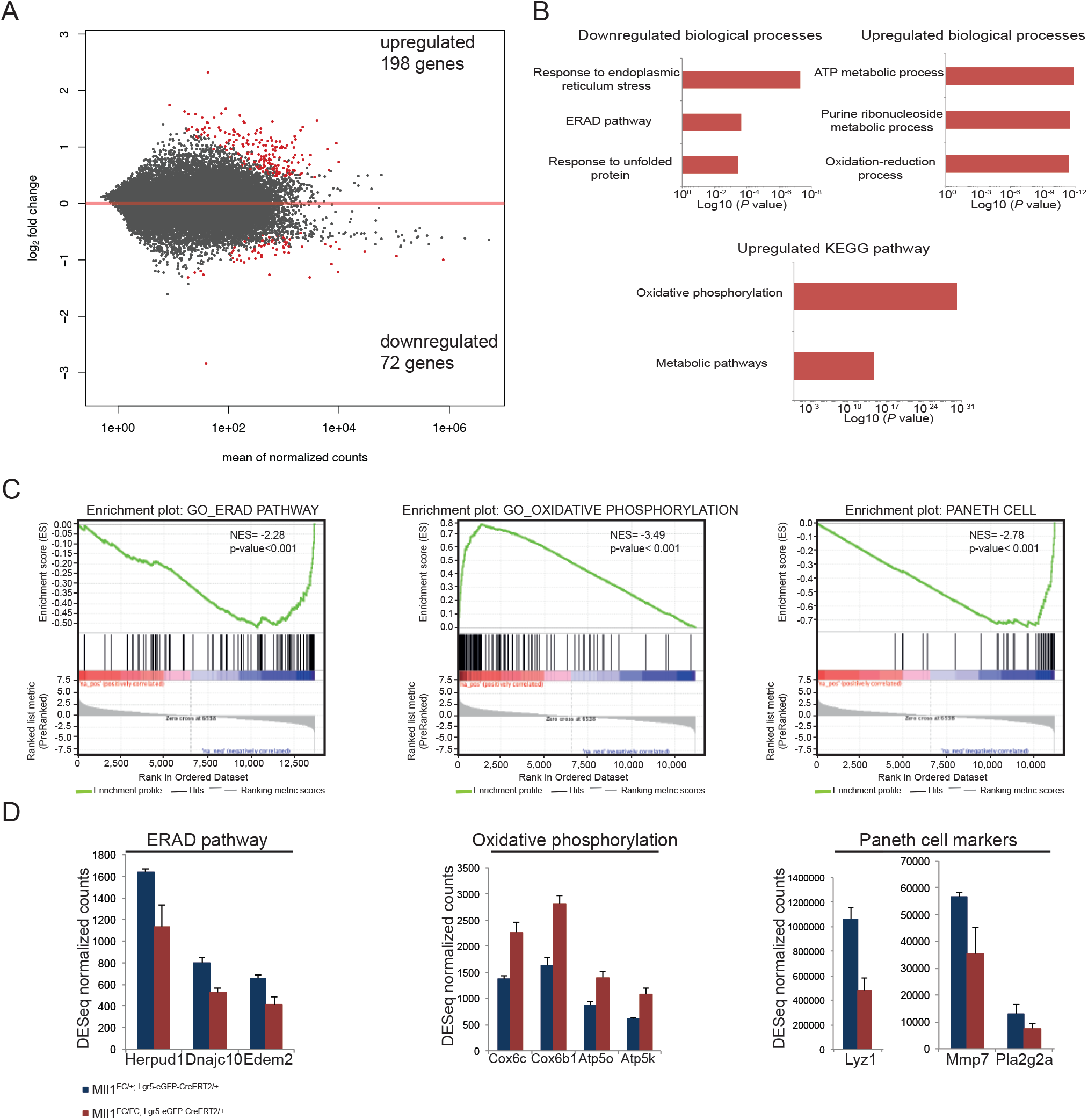
RNA profiling of wt Paneth cells after deletion of *Mll1* in the neighboring ISCs. (A) Wt Paneth cells neighboring either *Mll1*^*FC*/+; *Lgr5-eGFP-CreERT2*/+^ or *Mll1*^*FC/FC; Lgr5-eGFP-CreERT2*/+^ ISCs were sorted 4 days after tamoxifen induction was completed and were subjected to RNA profiling. MA plot visualizing the log2-fold change differences according to expression levels of Paneth cells. Red dots represent significant DEGs at a 5% FDR. (B) Enriched terms of biological processes and pathways down- and upregulated using DAVID GO/BP/FAT and KEGG database. (C) GSEA shows significant negative or positive correlation of genes from the GO ERAD pathway, GO oxidative phosphorylation and Paneth cell signature gene set in wt Paneth cells neighboring either *Mll1*^*FC/FC; Lgr5-eGFP-CreERT2*/+^ or *Mll1*^*FC*/+; *Lgr5-eGFP-CreERT2*/+^ ISCs. The Paneth cell signature gene set originates from [44]. NES: normalized enrichment score. (D) DESeq normalized counts for selected genes differentially regulated in the ERAD pathway, oxidative phosphorylation and Paneth cell signature gene set. Mean+s.d. is shown; n=4; p<0.05, Wald test.

### Skewed differentiation of organoids after loss of MLL1

To evaluate the cell-intrinsic requirement of MLL1 in the small intestine, we isolated crypts from *Mll1*^*F*/+; *RC*/+^ and *Mll1*^*F/F; RC*/+^ mice and cultured them to form organoids. After passaging, organoids were induced with 4-OH tamoxifen for 24 hours on day 2. After further passages, the *Mll1*^*FC/FC; RC*/+^ organoids increasingly formed round, less differentiated cyst-like spheres (Figs 6A-6C). qRT-PCR revealed mRNA downregulation of *Jaml* and transcription factors such as *Pitx2 and Foxa1* (Fig 6D) indicating the expected loss of ISCs and elevation of the goblet cell marker, *Muc2*. Notably the cyst-like organoids kept proliferating after loss of MLL1 and the Lgr5^+^ ISC marker, *Olfm4*, was elevated indicating differences between events in the crypt and in culture. Differences are also indicated by the elevation of the Paneth cell markers *Mmp7*, *Wnt3* and the elevation of the putative JAML receptor, *Cxadr*, which might enable the transition from organoids to spheroids (Fig 6D). These data establish a cell intrinsic requirement for MLL1 and indicate that it is required to maintain ISCs in organoid cultures.

**Fig 6.**
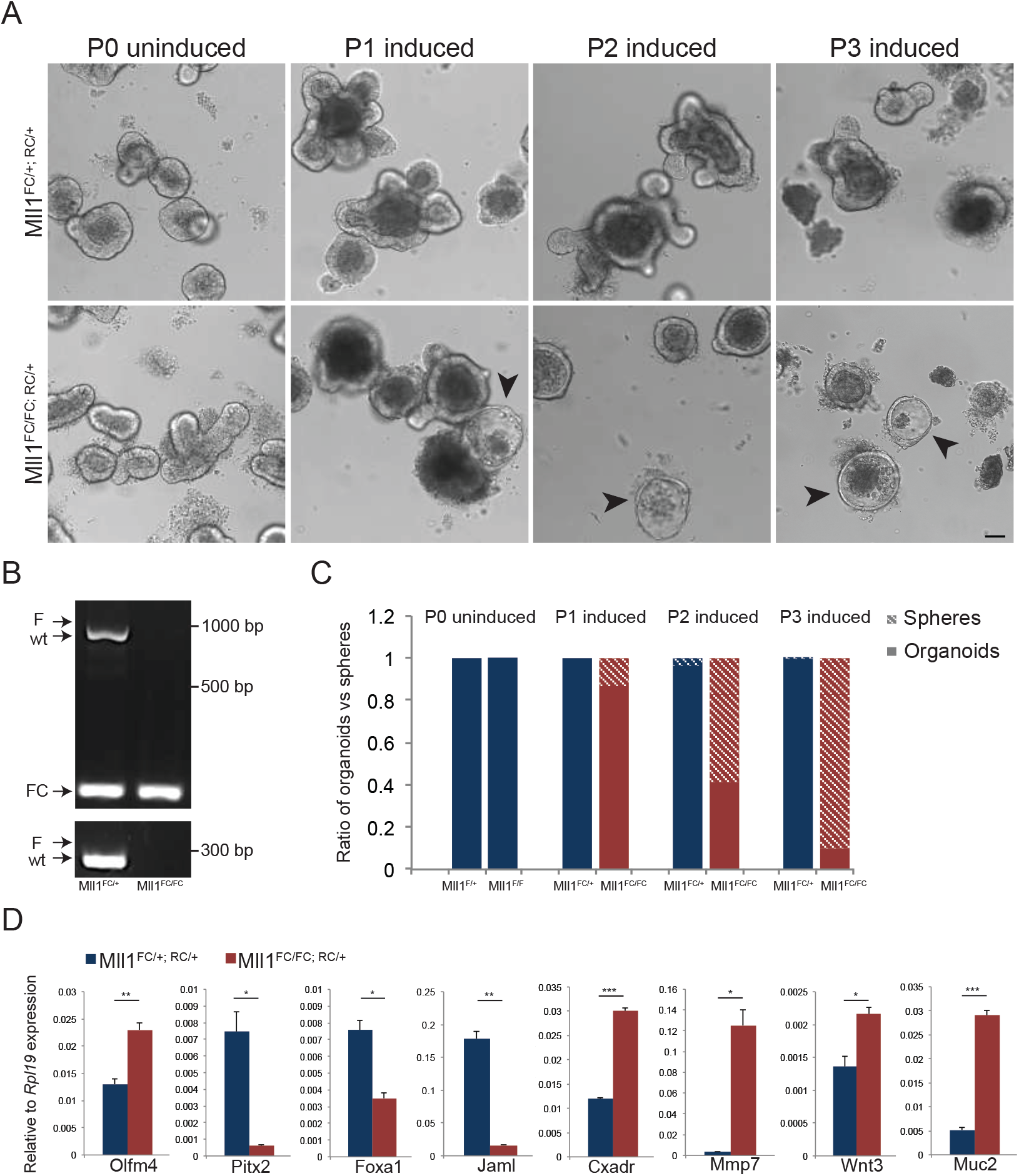
*Mll1* deletion in organoids results in formation of spheres. (A) Differential interference contrast (DIC) images of *Mll1*^*FC*/+; *RC*/+^ and *Mll1*^*FC/FC; RC*/+^ organoids. Organoids were induced with 4-OH tamoxifen for 24 h. Upon passaging (P), the mutant starts to loose its budding morphology giving rise to an undifferentiated cyst-like appearance. Scale bar 100 μm. (B) Genotyping of organoids/spheres for *Mll1* at passage 2. After tamoxifen induction the *Mll1^F^* allele recombines and results in the *Mll1^FC^* allele. PCR primers located upstream of the 5’ FRT site and downstream of the 3’ loxP site (see S1A Fig) identify the respective band. Consequently, the *Mll1^F^* band is 1084 bp, the wt band is 933 bp and the *Mll1^FC^* band is 186 bp. An additional PCR was performed with primers flanking the 3’ loxP site. Consequently, the *Mll1^F^* band is 297 bp and the wt band is 251 bp. (C) Quantification of organoids and spheres shown in A. (D) qRT-PCR was performed for selected genes on cDNA from *Mll1*^*FC*/+; *RC*/+^ and *Mll1*^*FC/FC; RC*/+^ organoid culture. Mean+s.d. is shown; n=3; *p<0.05, **p<0.01, ***p<0.001, Student’s *t* test.

## Discussion

Here we add a fourth stem cell to the known MLL1 repertoire of (i) HSCs [20, 21], (ii) skeletal muscle satellite cells [26] and (iii) postnatal neural stem cells (NSCs) [27]. The complete concordance between known MLL1 functions in postnatal stem cells suggests that MLL1 conveys an essential stem cell property. This possibility is enhanced by comparison to the MLL1 paralogue, MLL2, whose known functions in adult mice do not relate to stem cells rather macrophages (to respond to lipopolysaccharides) or fertility [48–51].

To explore the idea that MLL1 conveys a key stem cell property, we inspected the transcriptome profiles after conditional loss of MLL1 in the four adult/postnatal stem cells [26, 27, 52] (Supplementary excel file 1). However no shared candidate regulators or gene expression programs were identified. Although deeper, more systematic, transcriptome or cell biology approaches may reveal a shared MLL1 stem cell property, the lack of concordance between MLL1 regulation of these four stem cell transcriptomes is not unexpected. Previous work with MLL1 noted that direct target genes are not shared between different cell types [52] and a similar observation was made for MLL2 [48]. That is, the regulation of gene expression by the Trithorax homologues, MLL1 and MLL2, varies depending on the cell type and is not universal.

As again documented here for ISCs, the strongest relationship between the loss of MLL1 and cellular processes involves the downregulation of mRNAs that regulate transcription. Upon loss of MLL1, downregulation of transcription factor mRNAs include - (i) in HSCs; *Mecom, Prdm16, Pbx1, Eya1, Meis1* and *Hoxa9*; (ii) in postnatal NSCs; *Nkx2.1, Nkx2.3*; (iii) in satellite cells; *Pax7* and (iv) in ISCs; *Pitx2, Foxa1, Gata4* and *Onecut2*.

How does MLL1 regulate key lineage specific transcription factors differently in different lineages? MLL1 and MLL2 are amongst the few proteins that include the CxxC zinc finger that binds unmethylated CpG dinucleotides [23, 53] as well as a PHD finger that binds H3K4me3 [54]. Hence, as suggested before [48], MLL1 and 2 have the potential ability to bind CpG island promoters without the need for recruitment by sequence specific DNA binding transcription factors. This potential accords with the observation that both MLL1 and 2 appear to be bound at almost all active promoters [40, 55]. Consequently additional factors are required to explain the restricted transcriptional specificities of the MLLs. Notable in this regard, PAX7 is bound to MLL1 when satellite cells are activated, and enhanced transcriptional activation from both the *Myf5* promoter, to initiate skeletal muscle replenishment, and the *Pax7* promoter itself, depends on MLL1 [26]. This suggests that key transcription factors can either acquire the ability to interact with MLL1 bound at target promoters or recruit MLL1 to target promoters, or both.

Amongst the transcription factor mRNAs identified after loss of MLL1 in ISCs, *Pitx2* is prominent. *Pitx2* was previously identified as a direct target of MLL1 in ESCs and HSCs/hematopoietic progenitor cells [56, 57]. PITX2 is a homeodomain protein responsible for left-right asymmetric morphogenesis in the gut and proper positioning of the small intestine in the body cavity [58]. Also notably identified in ISCs are *Foxa1* and *Onecut2*. Both genes, previously known as *Hnf3α* and *Hnf6α*, are expressed in all epithelia of the gastrointestinal tract from its embryonic origin into adulthood. Together with *Math1*, they are critical for goblet cell differentiation and function [59, 60]. *Gata4*, which has previously been described as an MLL1 target gene [61], is also amongst the top downregulated mRNAs after loss of MLL1. Some aspects of the intestine specific deletion of *Gata4* in the adult mouse resemble the MLL1 phenotype described here including decreased proliferation in the crypts with increased numbers of goblet cells [62].

In addition to the central relationship between MLL1 and transcription factor expression, by far the most dramatically downregulated mRNA in both 4 and 10 day ISC transcriptomes was *Jaml*, which was previously included in the transcriptome profile that characterizes ISCs [43, 44]. As a prominent cell adhesion molecule, investigations by Tetteh and Clevers found that *Jaml* is expressed in the base of the crypt in ISCs but not Paneth cells and used *Villin-CreERT2* to conditionally knock it out [63]. Loss of *Jaml* resulted in loss of both *Olfm4*^+^ ISCs and proliferation in the crypt without loss of Paneth cells. These observations support the conclusion that MLL1 contributes to ISC function mainly by expression of *Jaml*. However in contrast to the loss of MLL1, Tetteh and Clevers did not observe an increase in goblet cells after loss of JAML. Imbalanced commitment in the secretory lineage may indicate a second aspect of MLL1 function that does not operate through *Jaml* expression, possibly including the regulation of *Gata4* expression [62] and other transcription factors. Upon loss of MLL1 *Jaml* is downregulated and the close association between stem cells and Paneth cells is probably destabilized (Fig 7). ISCs are anchored in the crypt surrounded by Paneth cells, which confers positional identity. The central ISC at the bottom of the crypt flanked by two Paneth cells has long-term self-renewal potential compared to border ISCs [64]. Interestingly positional identity of NSCs seems to be regulated in an MLL1 dependent fashion [27].

**Fig 7.**
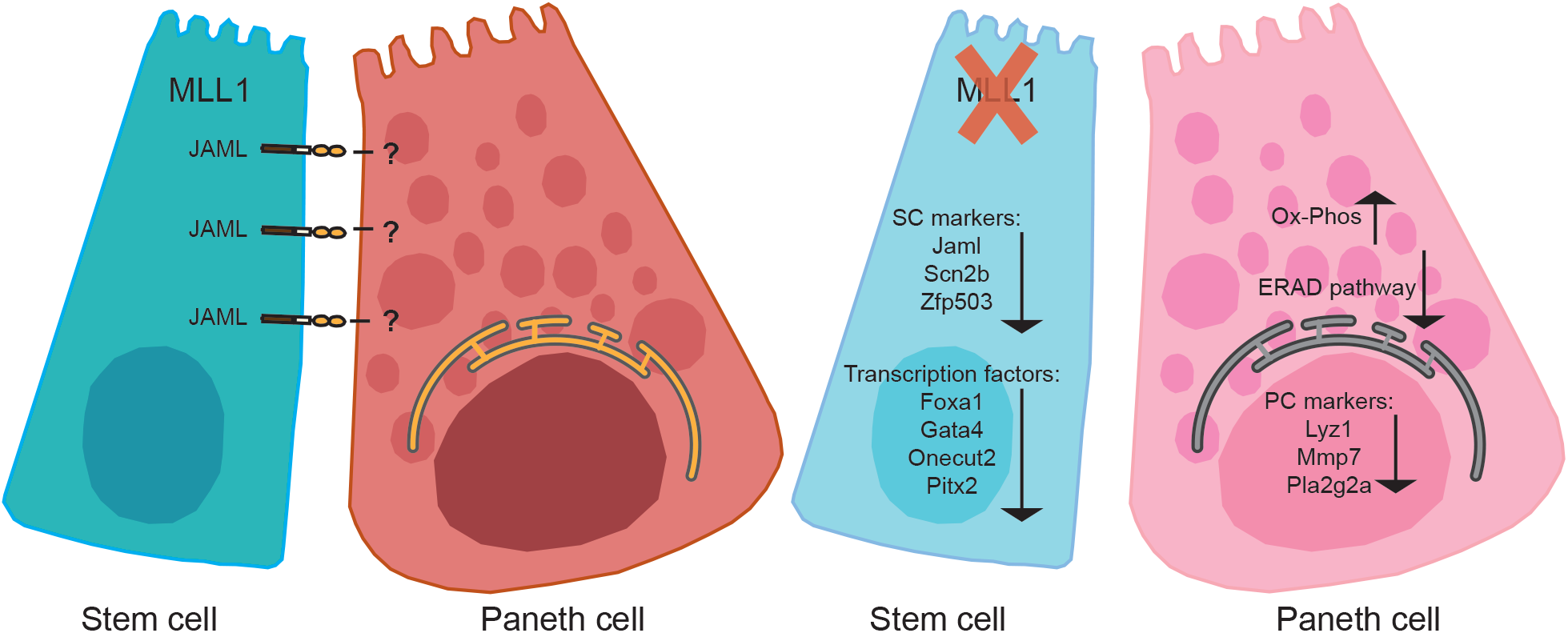
Cell-cell contacts are essential for cell identity. JAML is an integral transmembrane protein expressed on ISCs and interacts with unknown proteins on Paneth cells. Loss of MLL1 in ISCs causes loss of JAML mediated interactions, transcriptional changes and subsequently loss of stem and Paneth cell identity.

Once again we report the remarkably specific dependency of the expression of one or two genes on a Trithorax homologue. As described here, loss of MLL1 resulted in loss of *Jaml* expression with subsequent significant functional consequences in the crypt. In ESCs, the expression of one gene, *Magoh2*, entirely depends on MLL2. Removal of MLL2 resulted in the suppression of *Magoh2* expression by H3K27 methylation, followed by DNA methylation [48], thereby providing more evidence supporting the conclusion that a primary function of Trithorax action is to prevent Polycomb-Group repression [65]. These observations and conclusions with *Magoh2* in ESCs were recently confirmed [66]. Selective gene specific anti-repression could also explain the action of MLL2 on *Pigp* in macrophages [49] and MLL1 on *Hoxa9* in HSCs [25].

Despite a high degree of evolutionary conservation and near-ubiquitous expression, the Trithorax homologues appear to regulate a small number of genes in cell-type specific patterns, one or two of which are entirely dependent on one homologue for expression due to anti-repression. In ISCs this extraordinary specificity is focused on *Jaml* with functional consequences for the crypt niche. As observed in other adult stem cells, MLL1 also regulates the expression of transcription factors in ISCs that likely influence lineage commitment decisions. These observations lead to the attractive proposition that MLL1 is a master stem cell regulator. However a unifying molecular basis for this proposition remains to be identified.

## Methods and materials

### Targeting constructs

The targeting construct for *Mll1* was generated using recombineering (S1A Fig) employing an engrailed-intron-splice-acceptor-IRES-LacZ-Neomycin-polyA cassette flanked by FRT sites [39]. The critical exon 2, which upon deletion results in a frameshift and a premature stop codon in exon 3, was flanked by loxP sites.

### Gene targeting and generation of conditional knockout mice

Gene targeting in R1 embryonic stem cells (ESCs) was performed as described [67]. Correct integration in the *Mll1* locus was confirmed by Southern blot analysis using an internal probe and 5’ and 3’ external probes (S1B and S1C Figs). Two correctly targeted ESC clones were injected into blastocysts and gave rise to several chimeras, which were able to establish germ line transmission. *Mll1*^*A*/+^ mice were crossed to the *hACTB-Flpe* line to generate *Mll1*^*F*/+^ mice. *Mll1*^*A*/+^ and *Mll1*^*F*/+^ mice were backcrossed at least six generations to *C57BL/6JOlaHsd* mice. Subsequently, those mice were crossed to the *Rosa26-Cre-ERT2* (*RC*) line [5] to generate conditional, tamoxifen-inducible *Mll1*^*F*/+; *RC*/+^ mice. *Mll1*^*F*/+^ mice were bred with *Lgr5-eGFP-CreERT2* and *Villin-CreER^T2^* (*Vil-Cre-ERT2*) [29, 41] mice. Primers for genotyping are provided in S2 Table. Experiments were performed in accordance with German animal welfare legislation, and were approved by the relevant authorities, the Landesdirektion Dresden.

### Tamoxifen

Tamoxifen (Sigma Aldrich, T5648) was given to at least 10-week old mice by gavage (4.5 mg per day) for six days with three days break in between [68]. For RNA-Sequencing experiments, *Mll1*^*F*/+;^ and *Mll1*^*F/F; Lgr5-eGFP-CreERT2*/+^ mice received 1 mg tamoxifen via intraperitoneal (IP) injection for 3 consecutive days. Intestinal organoids were induced on day 2 after splitting using 800 nM 4-OH tamoxifen (Sigma H7904) for 24 h.

### Bone marrow transplantation

*Mll1*^*F*/+; *RC*/+^ and *Mll1*^*F/F; RC*/+^ recipient (CD45.2) mice were lethally irradiated with 8.5 Gy (X-ray source MaxiShot from Yxlon). Bone marrow cells from B6.SJL mice (CD45.1) were prepared by crushing with a mortar and pestle in ice-cold PBS supplemented with 5% fetal bovine serum. Red blood cells were removed with ACK lysis buffer (Thermo Fisher Scientific). 1 × 10^6^ lineage depleted (Lin^−^) bone marrow cells were injected into the retro-orbital venous plexus. Animals were maintained on water containing 1,17 mg/ml Neomycin (Merck) for three weeks after irradiation. Complete donor cell engraftment of wt CD45.1^+^ cells was confirmed by flow cytometry on peripheral blood with antibodies directed against the following murine antigens (clones given in brackets): CD45.1 (A20), CD45.2 (104), CD11b (M1/70), Gr-1 (RB6-8C5). Stably engrafted mice were fed six times with tamoxifen 30 weeks after transplantation. FACS analysis for KSL-Slam enriched HSCs was done with antibodies directed against the following murine antigens: CD3 (2C11; 17A2), CD11b (M1/70), CD16/32 (93), CD19 (eBio 1D3), CD34 (RAM34), CD45.1 (A20), CD45.2 (104), CD45R (RA3-6B2), CD117 (2B8), CD135 (A2F10), Gr-1 (RB6-8C5), Nk1.1 (PK136), Ter119 (Ter119, all eBioscience), CD11b (M1/70) and CD45.1 (A20, all BD Pharmingen), Sca-1 (D7), CD48 (HM48-1) and CD150 (TC15-12F1, all BioLegends). Lin^−^ cells were identified by lack of CD3, CD11b, CD19, CD45R, Gr-1, Nk1.1 and Ter119 expression.

### 5-Bromo-2-deoxyuridine (BrdU) assay

Mice were injected IP with BrdU (0.6 mg/10 g body weight in sterile PBS) and sacrificed 2 h later. The jejunum was dissected in cold PBS and processed for immunohistochemistry.

### Histochemistry and immunohistochemistry

Mouse intestine was flushed gently with cold PBS. Embryos were dissected from plugged mice on the respective gestational stage and placed in PBS. Intestine and embryos were fixed in 4% paraformaldehyde overnight. Dehydration and paraffin infiltration utilized the Paraffin-Infiltration-Processor (STP 420, Zeiss). Dehydrated tissues were embedded in paraffin (Paraffin Embedding Center EG1160, Leica) and 5 μm sections were prepared. Sections were deparaffinized in xylene, rehydrated through a series of alcohols, stained, dehydrated, and mounted. Femur sections were stained with Giemsa according to standard protocols. For the basic evaluation of the intestine hematoxylin and eosin (H&E) stain was performed. Periodic acid-Schiff (PAS) stain and alcian blue were used to identify goblet cells. For enterocytes, alkaline phosphatase stain was performed using two different methods. Sections were either incubated with Red Alkaline Phosphatase Substrate (Vector red, Vector Laboratories) for 10 minutes (min) or with nitroblue tetrazolium/5-bromo-4-chloro-3-indolyl phosphate solution for 30 min at room temperature (RT). For immunohistochemistry, antigen retrieval was performed by microwaving slides in 10 mM citrate buffer (pH 6.0) for 12 min (Microwave RHS 30, Diapath). Endogenous peroxidases were quenched with 0.3% H_2_O_2_ in methanol. Sections were incubated in blocking serum (5% goat serum) for 1 hr at RT followed by overnight incubation with primary antibodies (S3 Table) at 4°C. Following incubation with the secondary antibody (S3 Table) the immune peroxidase was detected using a Vectastain ELITE ABC kit (Vector) and visualized with a solution of diaminobenzidine (Sigma Aldrich) in the presence of 0.01% H_2_O_2_. All sections were counterstained with hematoxylin or alcian blue. Images were collected with an Olympus WF upright microscope and analyzed using the MetaMorph® Microscopy Automation and Image Analysis Software.

### *In situ* hybridization

The protocol for *in situ* hybridization was modified from [69]. Briefly, 8 μm thick sections were rehydrated. The sections were treated with 0.1 N HCl and proteinase K. Slides were postfixed and sections were then demethylated with acetic anhydride and prehybridized. Hybridization was done with 2 μg/ml digoxigenin (DIG)-labeled *Olfm4* RNA probe for 24 h at 65°C. Slides were washed and incubated with blocking solution for 1 hr. The sections were incubated with anti-DIG-alkaline phosphatase conjugate overnight at 4°C. Slides were washed and developed with BM purple.

### Crypt isolation

The jejunum was harvested from mice, flushed with ice cold PBS to remove any faecal content and cut open longitudinally. The tissue was placed lumen side up on a petri dish and the villi were removed by gently scraping the tissue using a glass cover slip. The tissue was cut into 2-4 cm pieces and was washed several times with ice-cold PBS to remove residual villi fragments. Tissues were transferred into a fresh tube containing 15 ml of 2 mM EDTA/PBS chelation buffer and placed on a rotating wheel for 30 min at 4°C.

The crypts were then detached from the basal membrane by vigorous shaking in 5% FCS/PBS solution. The suspension was filtered with a 100 μm cell strainer followed by a 70 μm cell strainer. Isolated crypts were centrifuged at 800 rpm for 5 min at 4°C. The final fraction consisted of pure crypts and was used for cell culture or single cell dissociation.

### Organoid culture

Purified crypts were resuspended in 10 ml DMEM/F12 (Life Technologies). 10 μl of the crypt suspension was used to count the number of crypts under the microscope. The pelleted crypts were resuspended in Matrigel^®^ matrix (Corning) at desired crypt density. Approximately, 400 crypts in 25 μl Matrigel^®^ matrix were seeded per well in a pre-warmed 24-well plate and incubated for 15 min at 37°C until the Matrigel^®^ matrix solidified. Then, 400 μl of IntestiCult™ Organoid Growth Medium (STEMCELL Technologies) was added to each well. Organoids were cultured at 37°C in a 5% CO_2_ incubator and maintained in culture for 5 days before being passaged and split for experimental procedures. The growth medium was replaced every 2-3 days

### Flow cytometry

ISCs were initially characterized and identified with the use of an *Lgr5-eGFP-CreERT2* knockin allele [29]. *Mll1*^*F*/+^ and *Mll1*^*F/F; Lgr5-eGFP-CreERT2*/+^ littermates (n=4) were given tamoxifen via IP injections for 3 consecutive days and were dissected 4 and 10 days later. Crypts were dissociated into single cells with TrypLE Express (Thermo Fisher Scientific) for 30 min at 37°C. Dissociated cells were passed through 70 μm cell strainer and washed with 5% FCS/PBS. Cells were stained with antibodies with the following antibodies for 45 min on ice: anti-mouse CD24 PE (clone M1/69), anti-mouse CD326 (EpCAM) APC (clone G8.8), anti-mouse CD45 Alexa-Fluor 700 (clone 104). Sorting was performed on a FACS Aria^TM^ III cell sorter (BD). After scatter discrimination to remove doublets the cell suspension was negatively selected with SYTOX blue dead cell stain and anti-CD45 to remove dead and hematopoietic cells, respectively. The cells where then positively selected with anti-EpCAM to enrich for intestinal epithelial cells. According to [70] CD45^−^, EpCAM^high^, CD24^med^ and GFP^high^ characterized ISCs and CD45^−^, EpCAM^high^, CD24^high^ and GFP^−^ characterized Paneth cells (S3A and S3B Figs). *Mll1* recombination via PCR on the sorted populations confirmed deletion of *Mll1* solely in the stem cell compartment (S3C Fig).

### RNA sequencing

300 intestinal stem and Paneth cells were sorted into 2 μl of nuclease free water with 0.2% Triton-X 100 and 4 U murine RNase Inhibitor (NEB). RNA was reverse transcribed (Invitrogen) and cDNA amplified using Kapa HiFi HotStart Readymix (Roche). The cDNA quality and concentration was determined with the Fragment Analyzer (Agilent). Samples were subjected to library preparation (TruePrep DNA library Prep Kit V2 for Illumina, Vazyme). Libraries were purified followed by Illumina sequencing on a Nextseq500 with a sample sequencing depth of 30 million reads on average. The short reads were aligned to the mm10 transcriptome with GSNAP (2018-07-04) and a table of read counts per gene was created based on the overlap of the uniquely mapped reads with the Ensembl Gene annotation version 92, using featureCounts (version 1.6.3). Normalization of the raw read counts based on the library size and testing for differential gene expression between the different genotypes was performed using the DESeq2 R package (version 1.24.0). Genes with an adjusted p-value (padj)≤ 0.05 were considered as significantly differentially expressed accepting a 5% FDR. To identify enrichment for particular biological processes and pathways associated with the DEGs, the DAVID GO/BP/FAT and KEGG database [71] was used. Gene set enrichment analysis was performed using GSEA software from the Broad Institute [72].

### Reverse transcription and quantitative PCR (qRT-PCR) analysis

RNA from sorted cells and organoids was extracted using Trizol (Sigma-Aldrich) and reverse transcribed using AffinityScript Multiple Temperature cDNA Synthesis Kit (Agilent Technologies). Real-time quantitative PCR was performed with GoTaq qPCR Master Mix (Promega) by Mx3000P QPCR System (Agilent Technologies). Ct values were normalized against *Rpl19*. Primer sequences and length of the amplified products are given in S2 Table. Fold differences in expression levels were calculated according to the 2^−ΔCt^ method [73].

### Quantification and Statistical analysis

Data is presented as mean and error bars indicate standard deviation (s.d.) unless otherwise indicated. Statistical details of the experiments can be found in the figure legends. Graphs and statistics were generated with GraphPad Prism software (v6.0) and Microsoft excel. Significance (p-values) for Kaplan-Meier graphs was determined by Mantel-Cox test. Significance (p-values < 0.05) was determined with Wald test or two-tailed Student’s *t* test. Significance for the breeding statistics was determined with Chi-square test. N indicates the numbers of independent biological replicates per experiment unless otherwise indicated.

## Data availability

RNA sequencing data have been deposited in the Gene Expression Omnibus under accession number GSE 157285.

## Acknowledgements

We thank Mandy Obst, Isabell Kolbe, Heike Petzold and Stefanie Weidlich for excellent technical assistance. We also thank the Biomedical Services (BMS) of the Max Planck Institute of Molecular Cell Biology and Genetics, Dresden for the excellent service and technical assistance. We are grateful to Prof. Sebastian Zeissig (CRTD, Dresden) for helpful advice. We thank the core facilities of the Biotechnology Center for providing assistance with flow cytometry (Katja Schneider). The Advanced Imaging Facility, a core facility of the CMCB Technology Platform at TU Dresden, http://biotp.tu-dresden.de/facilities/advanced-imaging/ assisted this research.

## Supporting information

**S1 Fig. *Mll1* gene targeting, embryonic phenotype and aspects of expression. (A)** Diagram of the *Mll1* gene with numbered exons and the multipurpose allele (*Mll1^A^*). This allele is converted to *Mll1^F^* upon FLP recombination. Cre recombination leads to excision of the frameshifting exon 2 generating the conditional mutant allele (*Mll1^FC^*). Genotyping primers are depicted for the downstream loxP site (loxP1 – loxP2) and for Flp recombination (Flp se – loxP2). SA = splice acceptor, IRES = internal ribosome entry site, pA = polyadenylation signal, lacZ-neo = β-galactosidase and neomycin resistance gene, * depicts premature stop codon. **(B)** Schematic representation of the Southern blot strategy. For identifying correct targeted events in the *Mll1* locus, Southern blot analysis employed 5’ (blue box) and 3’ (red box) probes. **(C)** Southern blot analysis using 5’ and 3’ external probes. **(D)** Dissected embryos from *Mll1*^*A*/+^ intercrosses at E12.5. *Mll1*^*A/A*^ embryos had a pale liver (marked by arrow). **(E)** Antibody staining (brown) shows that MLL1 is expressed in crypts and TA compartment of the small intestine but is absent in the villus (hematoxylin, purple). Scale bar 50 μm. **(F)** Normalized RNA-sequence counts for *Mll1/Kmt2a, Mll2/Kmt2b, Mll3/Kmt2c, Mll4/Kmt2d, Setd1a/Kmt2f and Setd1b/Kmt2g* in ISCs (eGFP^high^) and Paneth cells sorted from *Lgr5-eGFP-CreERT2* mice. Mean+s.d. is shown; n=4; *p<0.05, **p<0.01, ***p<0.001, ****p<0.0001, Student’s *t* test. **(G)** Antibody stainings of H3K4me1, H3K4me2 and H3K4me3 are comparable in *Mll1*^*FC*/+; *RC*/+^ and *Mll1*^*FC/FC; RC*/+^ intestinal sections. Scale bars are 100 μm.

**S2 Fig. Without bone marrow transplantation, *Mll1* deletion recapitulates the BMTx *Mll1* mutant phenotype. (A)** Antibody stain (left panels) and *in situ* hybridization (right panels) to visualize OLFM4/*Olfm4* in intestinal sections. Arrowheads point towards ISCs. Scale bar 100 μm. **(B)** Proliferative activity visualized by both Ki67 stain and BrdU incorporation in intestinal sections. Arrowheads point towards proliferative ISCs. Scale bars are 50 μm. **(C)** PAS staining and GOB5 antibody stain to visualize goblet cells in intestinal sections. Scale bars are 100 μm. **(D)** Chromogranin A and alkaline phosphatase staining to visualize enteroendocrine cells and enterocytes respectively. Arrows point to enteroendocrine cells (brown cytoplasmic stain) in the villi. Blue enterocytes covering the villi are marked by arrowheads. Scale bars are 100 μm for chromogranin A and 50 μm for alkaline phosphatase. **(E)** Nuclear β-catenin is comparable between the two different genotypes. Arrowheads point at β-catenin positive nuclei. Scale bar is 50 μm.

**S3 Fig. FACS gating strategy to sort ISCs and Paneth cells. Flow sorting on (A)** *Mll1*^*FC*/+; *Lgr5-eGFP-CreERT2*/+^ and **(B)** *Mll1*^*FC*/+; *Lgr5-eGFP-CreERT2*/+^ single cell suspension of crypts. Briefly, the consecutive gating steps were applied: (i) – (iii) Definition of the population of interest by exclusion of debris based on size (FSC), granularity (SSC) and the selection for single cells; (iv) Exclusion of dead cells that incorporated the nucleic acid stain SYTOX blue; (v) Depletion of CD45^pos^ population; (vi) Definition of Paneth (EpCAM^high^/CD24^high^) cell population by plotting EpCAM vs CD24 fluorescence; (vii) EpCAM^high^/CD24^med^ cell population was gated to discriminate the stem cell population (GFP^high^). **(C)** Stem cells (SC) and Paneth cells (PC) from *Mll1*^*FC*/+; *Lgr5-eGFP-CreERT2*/+^ and *Mll1*^*FC/FC; Lgr5-eGFP-CreERT2*/+^ mice 4 days after tamoxifen induction were checked for recombination. Left panel; PCR genotyping was using primers upstream of the 5’ FRT site and downstream of the 3’ loxP site identified the *Mll1^F^* band at 1084 bp, the wild type band a 933 bp and the *Mll1^FC^* band at 186 bp. Right panel; primers flanking the 3’ loxP site identified the *Mll1^F^* band at 297 bp and the wild type band at 251 bp.

**S4 Fig. Alignment and quality of the sequenced data. (A)** ISCs and Paneth cells were analyzed from control (*Mll1*^*FC*/+; *Lgr5-eGFP-CreERT2*/+^) (ctrl) (n=4) and *Mll1*^*FC/FC; Lgr5-eGFP-CreERT2*/+^ (n=4) (KO) mice. Mappability of reads for sorted ISCs 4 days after tamoxifen induction was completed. **(B)** Mappability of reads for sorted Paneth cells 4 days after tamoxifen induction was completed. **(C)** Mappability of reads for sorted ISCs 10 days after tamoxifen induction was completed. ISCs were analyzed from control (*Mll1*^*FC*/+; *Lgr5-eGFP-CreERT2*/+^) (ctrl) (n=4) and *Mll1*^*FC/FC; Lgr5-eGFP-CreERT2*/+^ (n=3) (KO) mice. **(D)** Principal-component analysis (PCA) was performed on Paneth cell and ISC samples sorted 4 days after tamoxifen. PCA is based on mRNA changes for the top 500 most diverse genes of stem cell (SC) and Paneth cell (PC) samples in comparison to published datasets for Lgr5^+^ SC [42] and CD24^+^ PC [35].

**S5 Fig. ISCs lacking MLL1 loose their cellular identity. (A) (B)** GSEA shows significant negative or positive correlation of genes from the stem (A) and goblet cell (B) signature gene set in *Mll1*^*FC/FC; Lgr5-eGFP-CreERT2*/+^ ISCs compared to control ISCs 10 days after tamoxifen induction was completed. The signature gene sets originate from [44]. NES: normalized enrichment score. **(C)** To validate RNA-seq results qRT-PCR was performed for selected genes on cDNA from *Mll1*^*FC*/+; *Lgr5-eGFP-CreERT2*/+^ and *Mll1*^*FC*/+; *Lgr5-eGFP-CreERT2*/+^ sorted stem cells 4 days after tamoxifen induction was completed. Mean+s.d. is shown; n=3; *p<0.05, **p<0.01, Student’s *t* test. **(D)** DESeq normalized counts demonstrate *Jaml* being highly expressed in Lgr5^+^ ISCs but not in Paneth cells. Mean+s.d. is shown; n=4; ***p<0.001, Student’s *t* test.

